# *dms-view*: Interactive visualization tool for deep mutational scanning data

**DOI:** 10.1101/2020.05.14.096842

**Authors:** Sarah K. Hilton, John Huddleston, Allison Black, Khrystyna North, Adam S. Dingens, Trevor Bedford, Jesse D. Bloom

## Abstract

The high-throughput technique of deep mutational scanning (DMS) has recently made it possible to experimentally measure the effects of all amino-acid mutations to a protein (Fowler and Fields 2014). Over the past five years, this technique has been used to study dozens of different proteins (Esposito et al. 2019) and answer a variety of research questions. For example, DMS has been used for protein engineering (Wrenbeck, Faber, and Whitehead 2017), understanding the human immune response to viruses (Lee et al. 2019), and interpreting human variation in a clinical setting (Starita et al. 2017; Gelman et al. 2019). Accompanying this proliferation of DMS studies has been the development of software tools (Bloom 2015; Rubin et al. 2017) and databases (Esposito et al. 2019) for data analysis and sharing. However, for many purposes it is important to also integrate and visualize the DMS data in the context of other information, such as the 3-D protein structure or natural sequence-variation data.

Here we describe *dms-view* (https://dms-view.github.io/), a flexible, web-based, interactive visualization tool for DMS data. *dms-view* is written in JavaScript and D3, and links site-level and mutation-level DMS data to a 3-D protein structure. The user can interactively select sites of interest to examine the DMS measurements in the context of the protein structure. *dms-view* tracks the input data and user selections in the URL, making it possible to save specific views of interactively generated visualizations to share with collaborators or to support a published study. Importantly, *dms-view* takes a flexible input data file so users can easily visualize their own DMS data in the context of protein structures of their choosing, and also incorporate additional information such amino-acid frequencies in natural alignments.

---

Users can access *dms-view* at https://dms-view.github.io. The tool consists of a data section at the top and a description section at the bottom. The data section displays the user-specified data in three panels: the site-plot panel, the mutation-plot panel, and the protein-structure panel (Figure 1A). When sites are selected in the site-plot panel, the individual mutation values are shown in the mutation-plot panel and highlighted on the protein structure. The user can toggle between site- and mutation-level metrics, which are defined in the user-generated input file. The description section is at the bottom of the page, and allows the user to add arbitrary notes that explain the experimental setup, acknowledge data sources, or provide other relevant information.

**Figure 1:**
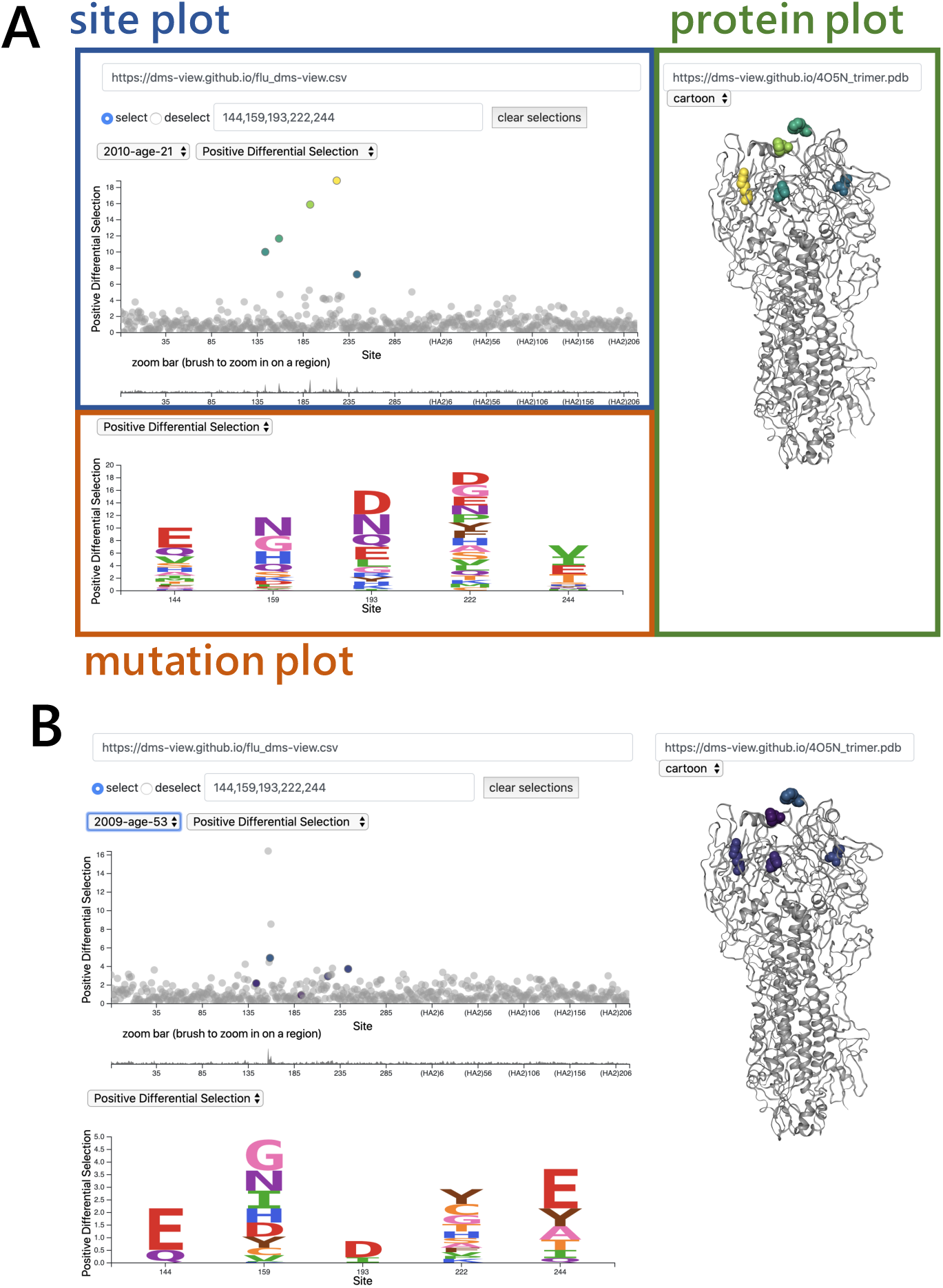
Using *dms-view* to analyze DMS data. For further exploration, please visit https://dms-view.github.io. **(A)** The *dms-view* data section has three panels: the site plot, the mutation plot, and the protein structure plot. The interactive features for selecting sites and navigating are in the site plot panel. Here we show the five sites most highly targeted by human serum “2010-Age-21” from the study by Lee et al. (2019). All five sites fall in the “globular head” of influenza virus HA. **(B)** The same five sites as in panel **A** but now plotted with the data from a different human serum, “2009-age-53”. Using *dms-view* to compare, we see that different sites on HA are targeted by different sera.

Please visit the documentation at https://dms-view.github.io/docs to learn more about how to use the tool, how to upload a new dataset, or view case studies.

## Example

### Mapping influenza A virus escape from human sera

Using a DMS approach, Lee et al. (2019) measured how all amino-acid mutations to the influenza virus surface-protein hemagglutinin (HA) affected viral neutralization by human sera. For more information on the experimental setup, see the paper (Lee et al. 2019) or the GitHub repo.

We visualized the Lee et al. (2019) serum mapping data using *dms-view*. To explore this dataset, please visit https://dms-view.github.io. In the *dms-view* visualization of these data, the conditions are the different human sera used for the selections. The site- and mutation-level metrics are different summary statistics measuring the extent that mutations escape from immune pressure.

Lee and colleagues asked two questions in their paper which can be easily explored using *dms-view*.

1. *Are the same sites selected by sera from different people?* To explore this question, we compared the site-level and mutation-level metric values for a specific set of sites between different conditions.
2. *Where on the protein structure are the highly selected sites located?* To explore this question, we selected specific sites of interest to be visualized on the 3-D protein structure

### Comparing site-level and mutation-level metric values for specific sites between conditions

To address whether or not the same sites are selected by different human sera using *dms-view*, we highlighted the most highly targeted sites for the human sera condition “Age 21 2010” Figure 1A (144, 159, 193, 222, and 244). We then used the condition dropdown menu to toggle to the other sera. The highlighted sites remain highlighted after the condition is changed so we can easily see if the same sites are targeted in other conditions.

In Figure 1B, we can see that there is no overlap of the sites selected by the human sera “2010-age-21” the human sera “2009-age-53”. These data are the default data for *dms-view*, so to explore this question in more detail please see https://dms-view.github.io.

### View sites on the protein structure

To address where on the protein structure the targeted sites are located, we selected the most highly targeted sites (144, 159, 193, and 222) for the human sera condition “Age 21 2010” to highlight them on the protein structure.

In Figure 1A, we can see that these sites cluster on the “head” of HA, which is known to be a common target of the human immune system (Chambers et al. (2015)).

## Code Availability

- dms-view is available at https://dms-view.github.io.
- Source code is available at https://github.com/dms-view/dms-view.github.io.
- Documentation (https://dms-view.github.io/docs) and case studies (https://dms-view.github.io/docs/casestudies/) are also available.

## Acknowledgements

This work started as the final project for UW class CSE 512 Data Visualization as a part of the UW eScience Advanced Data Science Option curriculum and we would like to thank Dr. Jeffrey Heer, Halden Lin, and Jane Hoffswell for their input on the initial design. Thank you to Bloom and Bedford lab members for their generosity providing feedback, data, and time for testing. This work was supported in part by the following grants of the NIAID of the NIH: F31AI140714, R01AI127983, R01AI141707, and R01AI140891. AB is supported by the National Science Foundation Graduate Research Fellowship Program under Grant No. DGE-1256082. TB is a Pew Biomedical Scholar and is supported by the additional NIH grants NIGMS R35 GM119774-01 and NIAID U19 AI117891-01. JDB is an Investigator of the Howard Hughes Medical Institute.

